# *Plasmodium* DEH is ER-localized and crucial for oocyst mitotic division during malaria transmission

**DOI:** 10.1101/2020.07.01.182774

**Authors:** David S. Guttery, Rajan Pandey, David J. P. Ferguson, Richard J. Wall, Declan Brady, Dinesh Gupta, Anthony A. Holder, Rita Tewari

**Affiliations:** School of Life Sciences, Queens Medical Centre, University of Nottingham, Nottingham, UK; The Leicester Cancer Research Centre, College of Life Sciences, University of Leicester, Leicester LE2 7LX, UK; Translational Bioinformatics Group, International Centre for Genetic Engineering and Biotechnology, Aruna Asaf Ali Marg, New Delhi-110067, India; Department of Biological and Medical Sciences, Faculty of Health and Life Science, Oxford Brookes University, Oxford, UK; Nuffield Department of Clinical Laboratory Science, University of Oxford, John Radcliffe Hospital, Oxford, UK; Wellcome Trust Centre for Anti-Infectives Research, School of Life Sciences, University of Dundee, Dundee DD1 5EH, UK; The Francis Crick Institute, London NW1 1AT, UK

**Keywords:** DEH, lipid metabolism, oocyst development, *Plasmodium*, sporogony

## Abstract

Cells use fatty acids (FAs) for membrane biosynthesis, energy storage and the generation of signaling molecules. 3-hydroxyacyl-CoA dehydratase – DEH – is a key component of very long chain FA (VLCFA) synthesis. Here, we further characterized in-depth the location and function of DEH, applying *in silico* analysis, live cell imaging, reverse genetics and ultrastructure analysis using the mouse malaria model *Plasmodium berghei*. DEH is evolutionarily conserved across eukaryotes, with a single DEH in *Plasmodium* spp. and up to three orthologs in the other eukaryotes studied. DEH-GFP live-cell imaging showed strong GFP fluorescence throughout the life-cycle, with areas of localized expression in the cytoplasm and a circular ring pattern around the nucleus that colocalized with ER markers. Δ*deh* mutants showed a small but significant reduction in oocyst size compared to WT controls from day 10 post-infection onwards and endomitotic cell division and sporogony were completely ablated, blocking parasite transmission from mosquito to vertebrate host. Ultrastructure analysis confirmed degeneration of Δ*deh* oocysts, and a complete lack of sporozoite budding. Overall, DEH is evolutionarily conserved, localizes to the ER and plays a crucial role in sporogony.

## Introduction

Malaria remains one of the world’s deadliest infectious diseases. Caused by apicomplexan parasites belonging to genus *Plasmodium*, malaria is responsible for great socio-economic loss to affected countries. According to WHO reports, there were 212 million clinical cases of malaria infection and 429000 deaths in 2015 (WHO, 2018), and growing resistance against existing drugs has further intensified this problem. Hence, there is a growing need to identify new biological pathways and proteins essential for parasitic growth and development in both human and non-human hosts, which could act as suitable drug targets. *Plasmodium* parasites have a complex life cycle and require two hosts to complete the life cycle: vertebrates (during asexual stages) and invertebrates (during sexual stages) (Aly *et al.*, 2009). The disease is transmitted to vertebrate hosts by infected female *Anopheles* mosquitoes, which inject sporozoites into the dermis of the vertebrate host during a blood meal. The parasite enters the circulation, and once it invades liver cells, and subsequently erythrocytes, undergoes several rounds of atypical closed mitotic cell division through multiple rounds of DNA replication and asynchronous nuclear division (termed schizogony) to produce merozoites that invade erythrocytes. During this period of cyclic asexual proliferation in the blood stream, a subpopulation undergoes gametocytogenesis to develop into male and female gametocytes, which are transmitted to a mosquito during its blood meal. Gamete development, fertilization and zygote formation occur in the mosquito midgut, leading to the differentiation of an infective ookinete, which undergoes meiosis, penetrates the midgut wall and develops into an oocyst on the basal surface of the midgut, where further rounds of closed mitotic cell division occur. Thousands of sporozoites develop within each oocyst, and then egress into the haemocoel to invade the salivary glands and begin a new life cycle.

Lipid metabolism includes essential cellular processes that use fatty acids (FAs) in membrane biosynthesis, energy storage and the generation of signaling molecules. FA synthesis is a four-step cyclic process that results in the addition of two carbons to the chain with each cycle. In humans, the process involves condensation of acyl-CoA with malonyl-CoA to produce 3-ketoacyl-CoA (catalyzed by one of seven FA elongases), reduction of 3-ketoacyl-CoA by a 3-ketoacyl-CoA reductase (KAR) to 3-hydroxyacyl-CoA, dehydration of 3-hydroxyacyl-CoA to 2,3-trans-enoyl-CoA (catalysed by one of four 3-hydroxyacyl-CoA dehydratase isoenzymes; HACD1-4 also known as DEH in some systems [see below]), and finally reduction to an acyl-CoA with two additional carbon chain units by 2,3-trans-enoyl-CoA reductase (TER) (Kihara, 2012). HACD1-4 were initially annotated as PTPLA, PTPLB, PTPLAD1 and PTPLAD2, respectively due to their similarities to the yeast Phs1 gene product (Ikeda *et al.*, 2008). HACD1 and HACD2 genes restored growth of yeast SAY32 Phs1-defective cells, indicating that they are functional homologues of Phs1, i.e. 3‐hydroxyacyl‐CoA dehydratases. Further, studies have indicated that HACD1 has an essential role in myoblast proliferation and differentiation (Lin *et al.*, 2012), with HACD1-deficient cell lines displaying S-phase arrest, compromised G2/M transition and retarded cell growth. Studies of the *Arabidopsis thaliana* Phs1 homologue PASTICCINO2 or PAS2 showed the protein has an essential role in very long chain fatty acid (VLCFA) synthesis (Bach *et al.*, 2008), as well as being essential during cell division, proliferation and differentiation (Bellec *et al.*, 2002). Further, *Arabidopsis* PAS2 complements Phs1 function in a yeast mutant defective for FA elongation (Morineau *et al.*, 2016). PAS2 interacted with FA elongase subunits in the endoplasmic reticulum (ER) and in its absence 3-hydroxyacyl-CoA accumulates, as expected from loss of a dehydratase involved in FA elongation. Similarly, in the yeast *Saccharomyces cerevisiae* VLCFA synthesis is also catalyzed in the ER by a multi-protein elongase complex, following a similar reaction pathway as mitochondrial or cytosolic fatty acid synthesis (Tehlivets *et al.*, 2007).

In Apicomplexans, the process of fatty acid synthesis and assembly into more complex molecules is critical for their growth and development, while also determining their ability to colonize the host and to cause disease. They acquire lipids through *de novo* synthesis and through scavenging from the host (Mazumdar *et al.*, 2007), and simple components like mosquito-derived lipids determine within-host *Plasmodium* virulence by shaping sporogony and metabolic activity, affecting the quantity and quality of sporozoites, respectively (Costa *et al.*, 2018). Fatty acid synthesis (FAS) occurs in the apicoplast via the type II FAS (FASII) pathway, followed by fatty acid elongation (FAE) on the cytoplasmic face of the ER through the elongase (ELO) pathway (Ramakrishnan *et al.*, 2012, Ramakrishnan *et al.*, 2013) (Figure S1). Studies on whether FAS is essential suggest that different *Plasmodium* spp. have different requirements for these enzymes. In *Plasmodium yoelii* the FASII enzymes are only essential during liver stages (Yu *et al.*, 2008, Vaughan *et al.*, 2009); whereas in *Plasmodium falciparum*, genetic disruption of the FASII enzymes FabI and FabB/F results in complete abolition of sporogony (van Schaijk *et al.*, 2014). Specifically, day 17 to day 23 after mosquito feeding, FabB/F mutant oocysts appeared to degenerate, and protein expressed from the *dhfr* resistance marker fused with *gfp* in PfΔ*fabB/F* deletion mutants was barely detectable using fluorescence microscopy. The enzymatic steps of the ELO process are similar to those in the FASII pathway in the apicoplast (Tarun *et al.*, 2009); however, the growing chain is held by CoA instead of acyl carrier protein (ACP). The ELO pathway consists of 3 additional enzymes involved in condensation: ELO-A, ELO-B and ELO-C (Ramakrishnan *et al.*, 2012), with ELO-A and ELO-B engaged in the elongation of *de novo* synthesized unsaturated fatty acids and ELO-C primarily acting on host-derived saturated fatty acids. In *Toxoplasma gondii*, the activity of the ELO-pathway is considered an alternative route to FASII-independent ^14^C-acetate incorporation (Bisanz *et al.*, 2006) and is engaged in conventional elongation rather than *de novo* synthesis (Mazumdar *et al.*, 2007). Indeed, *P. falciparum* parasites with no functional FASII pathway can still elongate fatty acids; possibly because of the activity of the ELO pathway (Yu *et al.*, 2008). Genetic deletion studies of the ELO enzymes in *Toxoplasma* have suggested functional redundancy (Ramakrishnan *et al.*, 2012); whereas in *Plasmodium* ELO-A has a crucial role during liver-stage development (Stanway *et al.*, 2019).

In a genome-wide study of *Plasmodium berghei* (Pb) protein phosphatases, we identified 30 phosphatase genes together with one for a predicted protein tyrosine phosphatase-like protein, PbPTPLA, which was shown to be essential for sporozoite formation and completion of the parasite life cycle, but not fully characterized (Guttery *et al.*, 2014). However, despite the original annotation as an inactive PTP-like protein (Andreeva *et al.*, 2008, Wilkes *et al.*, 2008, Guttery *et al.*, 2014, Pandey *et al.*, 2014), more recent functional studies indicate that it is a key component of the VLCFA elongation cycle – more specifically the ELO pathway as a 3-hydroxyacyl-CoA dehydratase (DEH) (Stanway *et al.*, 2019). Therefore, to investigate further the role of DEH in *Plasmodium* development, we performed an in-depth genotypic and phenotypic analysis of the protein, using *in silico*, genetic manipulation and cell biological techniques. We show evolutionary conservation of DEH in the model organisms examined here. Furthermore, we show that PbDEH is located at the ER and is essential for cell division and parasite budding within oocysts, with its deletion blocking parasite transmission.

## Results

### Phylogenetic analysis reveals that DEH is highly conserved among eukaryotes

Genome-wide analysis showed DEH is present in all the eukaryotic organisms studied here, which includes apicomplexans, fungi, plants, nematodes, insects, birds and mammals. The number of encoded DEH proteins was shown to vary from one to three in the studied organisms, with *Plasmodium* spp genomes coding for a single DEH. Both *Arabidopsis thaliana* and *Oryza sativa* encode three DEHs (*PAS2* and 2 *HACD* isozymes) each, as compared to two (HACD1 and HACD2) in *Homo sapiens* (plus two sharing relatively weak similarity – HACD3 and HACD4) and two in *Mus musculus*. Phylogenetic analysis using the neighbor joining method, clustered organisms based on their evolutionary relatedness (Figure S2, Table S1). In addition, the phylogenetic analysis suggests that gene duplication in non-chordata, chordata and plants where there are multiple DEH copies, may have happened independently from a single DEH gene to perform specific functions after divergence during evolution, based on the grouping of all DEH isoforms in the same cluster.

### *Plasmodium* DEH does not contain the canonical PTPLA CXXGXXP motif and is predicted *in silico* to interact with factors associated with FAE

*P. berghei* (PBANKA_1346500) and *P. falciparum* (Pf; PF3D7_1331600) DEH genes are annotated as PTPLA (pfam04387) (Andreeva *et al.*, 2008, Wilkes *et al.*, 2008, Guttery *et al.*, 2014, Pandey *et al.*, 2014), the criterion for PTPLA being the presence of a PTP active site motif (CXXGXXR) but with arginine replaced by proline (CXXGXXP). However, CLUSTALW alignment of *Pb* and *Pf* protein sequences with the human and mouse HACD1 and HACD2 shows this motif is absent (Figure S3), indicating that *Plasmodium* DEHs cannot be classified as PTP-like proteins. Secondary structure prediction of high confidence showed the presence of six hydrophobic helices followed by coils, and the absence of beta sheets (Figure S4A). In the absence of a PbDEH crystal structure or a significant template for homology-based 3D modeling revealed by BLASTP, we used I-TASSER (an *ab initio* threading based tool) to predict the 3D structure of PbDEH. This analysis provided a prediction consistent with the secondary structure and of a three dimensional structural model with six membrane-spanning helices (Figure S4B).

STRING database analysis predicted that PfDEH interacts with FAE and FAS proteins, and many other proteins with an ER location (Figure S5). These proteins include 3-oxo-5-α-steroid 4-dehydrogenase (PBANKA_09127; PF3D7_1135900), Stearoyl-CoA δ-9 desaturase (PBANKA_1110700; PF3D7_0511200), putative long chain polyunsaturated fatty acid elongation enzyme (ELO-B, PBANKA_0104700; PF3D7_0605900 - involved in the FAE pathway) as well as β-hydroxyacyl-(Acyl-carrier-protein) dehydratase (FabZ), involved in stage 3 of fatty acid synthesis in the FASII pathway (Stanway *et al.*, 2019). In addition, interactome analysis revealed DEH interaction with a putative ER membrane protein, Acetyl-CoA transporter protein (PF3D7_1036800).

### DEH is expressed throughout the *Plasmodium* life-cycle stages and localized to the ER

To determine the expression profile and location of PbDEH, we used a single homologous recombination strategy to tag the 3’ end of the endogenous *deh* locus with sequence coding for GFP (Guttery *et al.*, 2014), and then analyzed blood and mosquito stages of the life-cycle for GFP. Strong GFP fluorescence was observed throughout all life-cycle stages analyzed, with areas of localized expression in the cytoplasm and a circular ring formation around the nucleus (Figure 1). Predotar analysis (Small *et al.*, 2004) predicted an ER localization for both PbDEH and PfDEH. Colocalization with ER tracker confirmed the DEH-GFP location at the ER, in all parasite stages analyzed (Figure 2A), with subcellular fractionation of blood stage parasites confirming its integral membrane location (Figure 2B).

**Figure 1:**
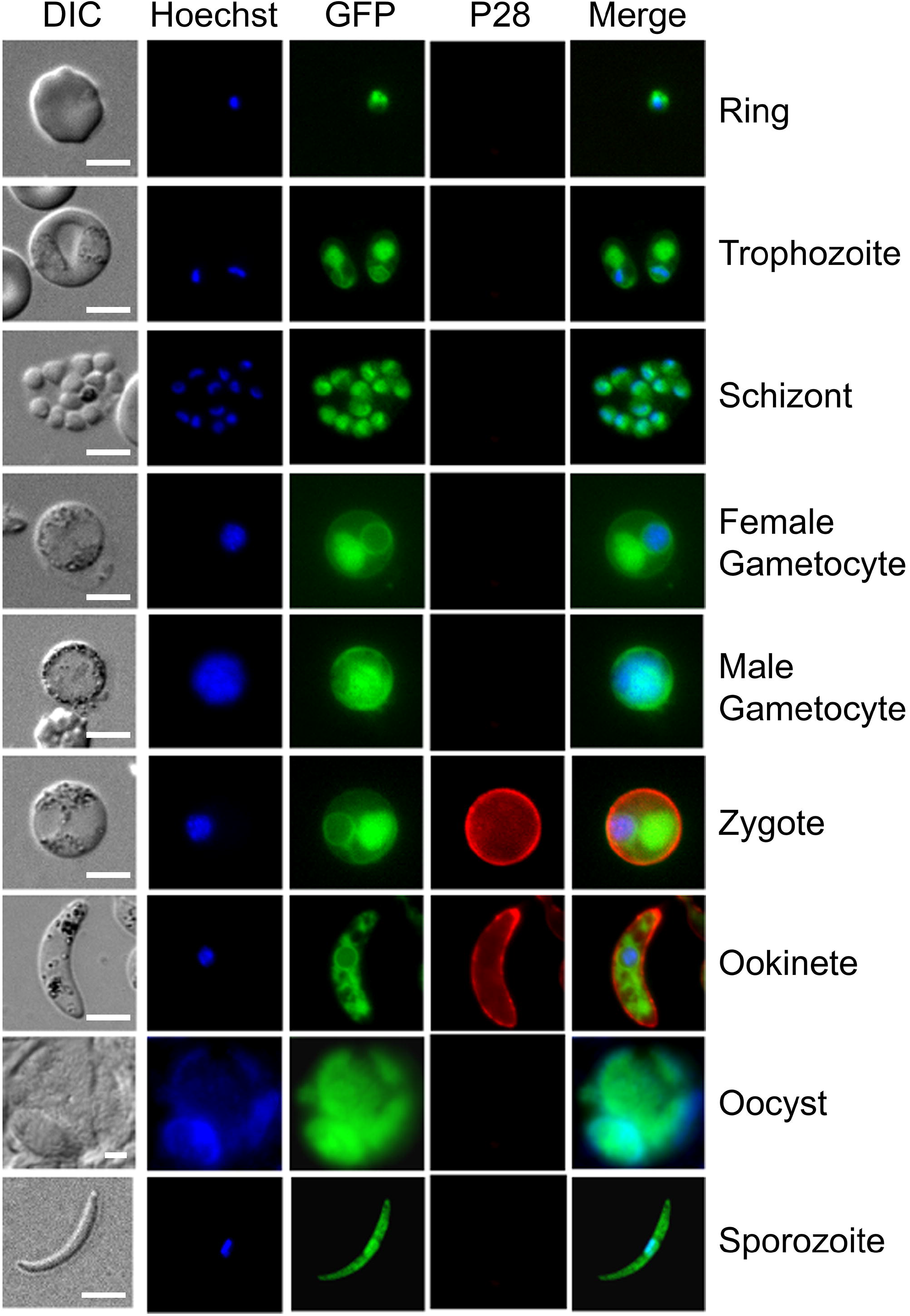
DEH-GFP protein expression in stages of the parasite life cycle. Expression of DEH-GFP in rings, trophozoites, schizonts, gametocytes, zygotes, ookinetes, oocysts and sporozoites. P28, a cy3-conjugated antibody which recognizes P28 on the surface of zygotes, and ookinetes was used as a marker of the sexual stages. Note that the female gametocyte in this figure has not been activated, and is not expressing P28. Scale bar = 5 μm.

**Figure 2:**
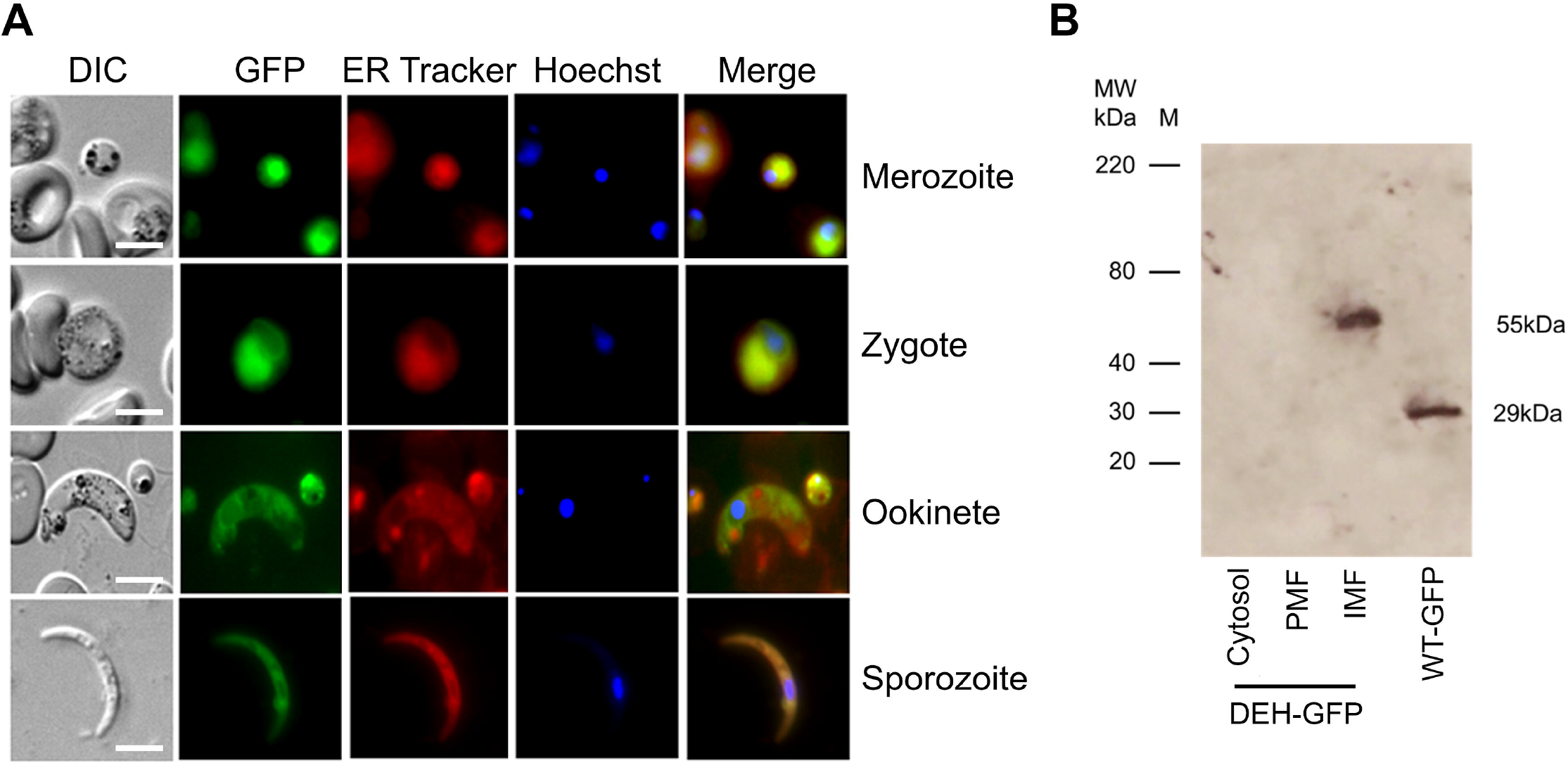
Co-localization of DEH-GFP and ER tracker. (A) Analysis of DEH-GFP localization using ER tracker in merozoites, zygotes, ookinetes and sporozoites. Scale bar = 5 μm. (B) Anti-GFP western blot for subcellular localization of DEH-GFP. PMF – Peripheral membrane protein fraction, IMF – Integral membrane protein fraction. For WT-GFP, cytosolic GFP is shown.

### PbDEH is essential for mitotic cell division of *Plasmodium* during oocyst development

Previous comparison of Δ*deh* and WT parasite lines highlighted the non-essential role of this gene for blood stage development (Guttery *et al.*, 2014). In this study we confirmed it is not essential during asexual blood stages, or for zygote development (Figure 3A). However, while the overall number of oocysts observed in Δ*deh* and WT lines was not significantly different (Guttery *et al.*, 2014), there was a significant reduction in Δ*deh* GFP-expressing oocysts beginning at day 7 and continuing through day 21 post-infection (Figure 3B, Table S2), with many appearing to be degenerating. Analysis of oocyst size revealed a small (but statistically significant) decrease from day 10 onwards in Δ*deh* lines compared to WT (Figure 3C, Table S2), and by day 21 the vast majority of Δ*deh* oocysts expressed GFP no longer, and in the few that did GFP was present at very low levels or in fragmented patterns. However, it is important to note that Δ*deh* oocysts that continued to express GFP and showed faint DAPI staining of DNA, were similar in size to WT oocysts; whereas the vast majority of oocysts that reduced in size did not express GFP or stain with DAPI, suggesting they were dead. Analysis of salivary glands from mosquitoes infected with Δ*deh* parasites revealed no sporozoites, in contrast with WT-parasite infected mosquitoes (Figure 3D). Representative examples of Δ*deh* oocyst morphology and lack of sporozoite development at all stages post-infection are shown in Figure 3E and Figure S6, highlighting oocyst degeneration, fragmented GFP expression and failure to form sporozoites.

**Figure 3:**
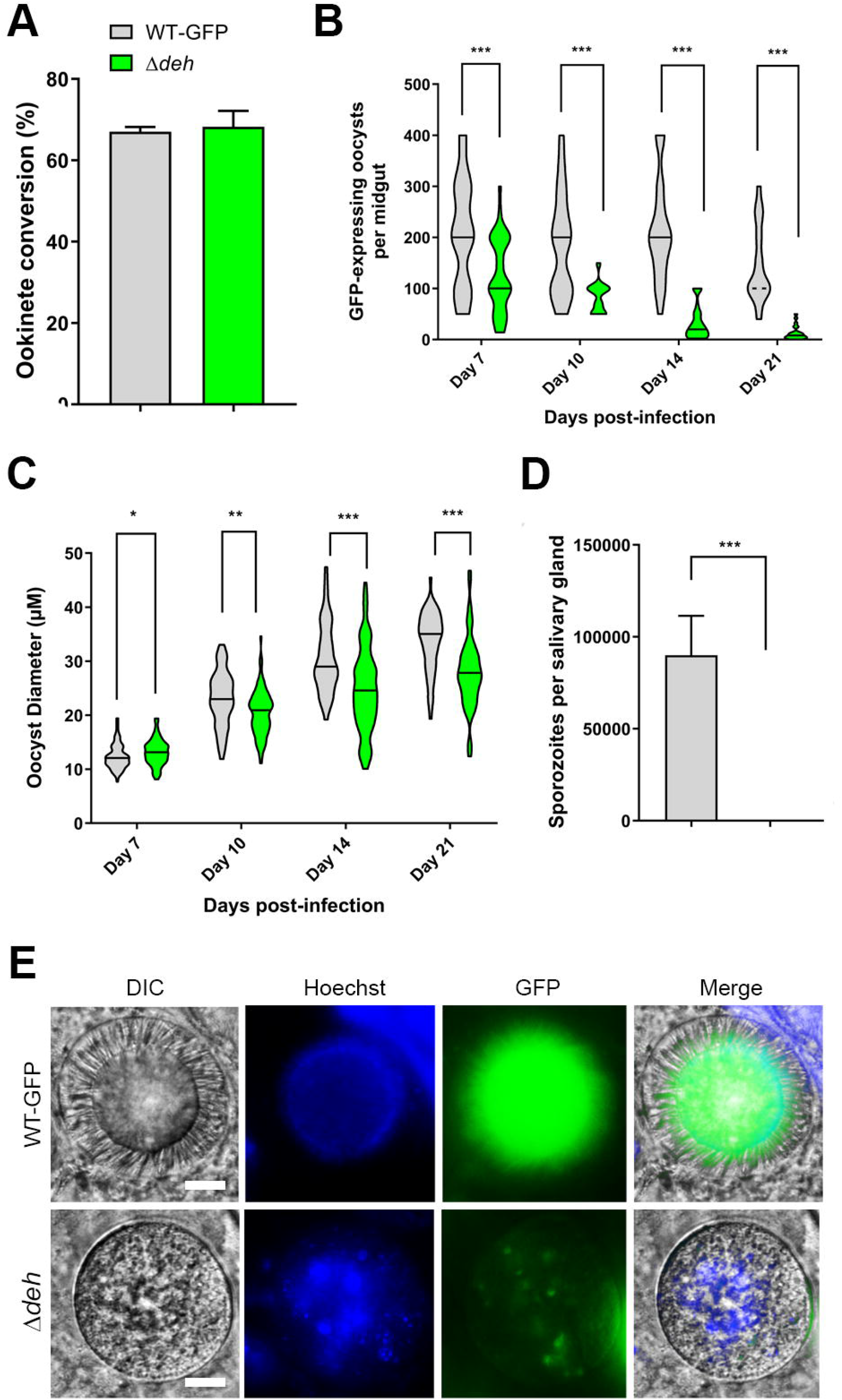
Phenotypic analysis of Δ*deh* lines. (A) Ookinete conversion as a percentage in Δ*deh* and WT lines. Ookinetes were identified using the marker P28 and defined as those cells that successfully differentiated into elongated ‘banana shaped’ ookinetes. Bar is the mean ± SEM. n = 3 independent experiments. (B) Total number of GFP-positive oocysts per infected mosquito, including normal and smaller oocysts, at 7, 10, 14, and 21 days post-infection for Δ*deh* and WT parasite lines. Bar is the mean ± SEM. n = 3 independent experiments (20 mosquitoes for each). P <0.001 for all time points. (C) Oocysts (10x magnification) of Δ*deh* and WT lines at 7, 10, 14 and 21 days post-infection. Scale bar = 100 μm. (D) Individual Δ*deh* and WT oocyst diameters (μm) at 7, 10, 14 and 21 days post-infection. Horizontal line indicates the mean from 3 independent experiments (20 mosquitoes for each) of Δ*deh* and WT. *p <0.05, **p <0.01, ***p <0.001. (E) Total number of sporozoites per mosquito from 21 days post-infection salivary glands for Δ*deh* and WT lines. Three independent experiments, n = 20 mosquitoes for each replicate. *** p<0.001. (F) Representative examples of Δ*deh* and WT oocysts (63x magnification) at 21 dpi showing fragmented GFP and Hoechst staining. Scale bar = 20 μm.

### Ultrastructure analysis confirmed oocyst degeneration and apoptotic-like nuclear chromatin condensation in Δ*deh* lines

To investigate further the marked differences in oocyst morphology and complete lack of sporozoite formation, we used electron microscopy to compare Δ*deh* and WT lines at 10-, 14- and 21-days post-infection. At 10 days, the oocysts of Δ*deh* and WT parasites were of a similar size. However, Δ*deh* oocysts showed numerous cytoplasmic vacuoles (Figure 4a) with evidence of dilatation of the nuclear membranes (Fig 4b). At day 14 post-infection the majority of Δ*deh* oocysts exhibited mid-(30%) or advanced (70%) stages of degeneration with increased cytoplasmic vacuolation, dilatated nuclear membranes, and evidence of mitochondrial abnormalities (Fig 4c, d). There was little evidence of retraction of the plasmalemma from the oocyst wall and no evidence that sporozoite inner membrane complex (IMC) formation had been initiated in any of the 20 oocysts examined. At day 21 post-infection, all Δ*deh* oocysts were in an advanced stage of degeneration - almost completely vacuolated with a few nuclei appearing to have undergone apoptotic-like nuclear chromatin condensation (Fig 4e, f). In contrast, at day 10 post-infection WT oocysts were completely filled with cytoplasm with numerous nuclear and mitochondrial profiles (Figure 4g, h). At 14 days post-infection the WT parasites showed a mixture of early (15%), mid (60%) and late (25%) stage oocysts with sporozoites at various stages of development (Figure 4i, j). At day 21 post-infection, the majority of WT oocysts (85%) were late stage with numerous fully formed and free sporozoites (Fig 4k, l) although a few mid-stage (10%) and rare (<5%) degenerate oocysts were observed.

**Figure 4:**
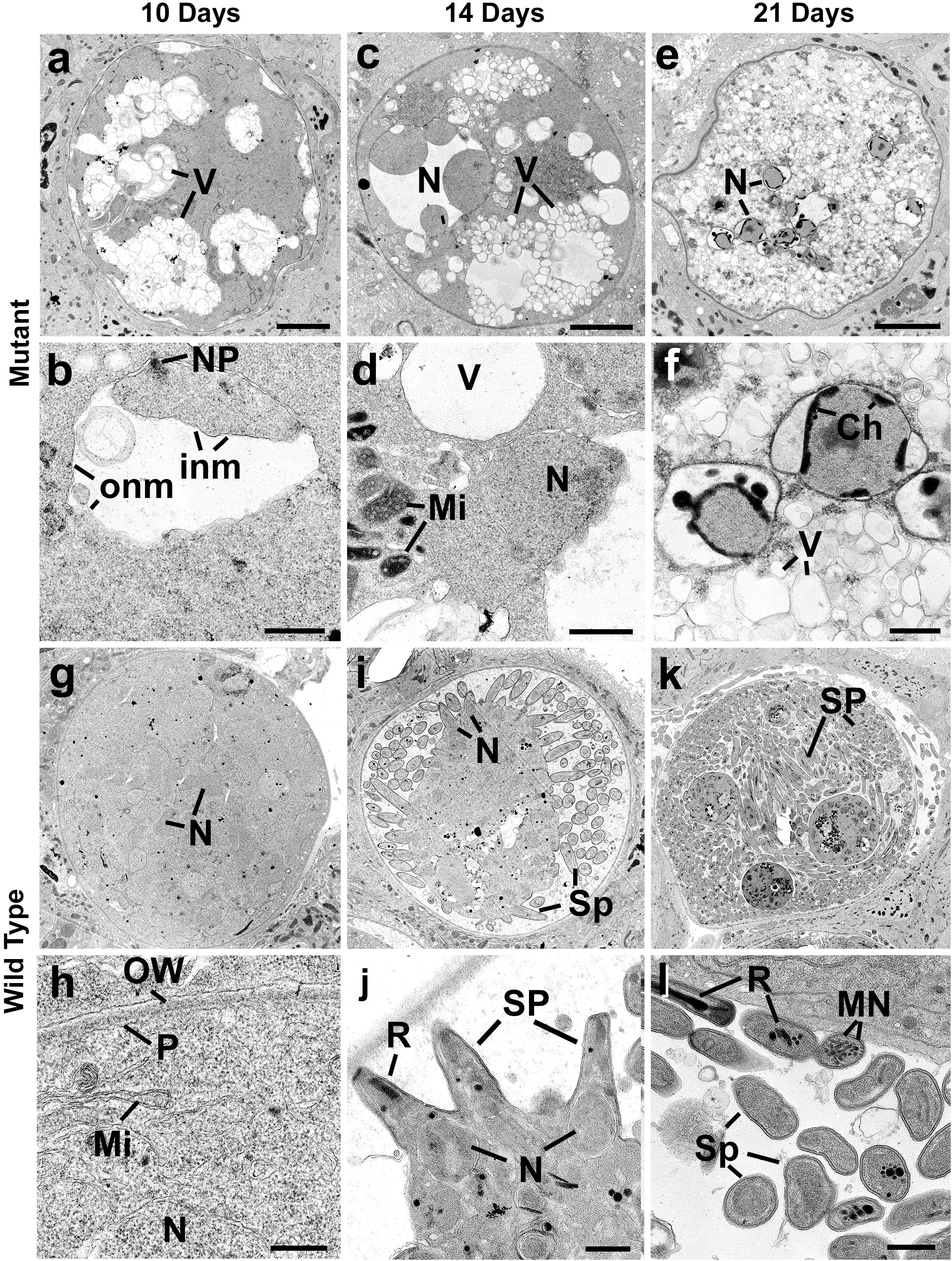
Ultrastructure analysis of oocyst development in Δ*deh* lines. Electron micrographs of Δ*deh* (a-f) and WT (g-l) parasites at 10 days (a, b, g, h), 14 days (c, d, i, j) and 21 days (e, f, k, l) post infection. Bars represent 10 μm (a, c, e, g, i, k) and 1 μm (b, d, f, h, j, l). (a) Low power image of an early oocyst showing lucent area made up numerous vacuoles (V). (b) Detail from a similar stage to that in (a) showing part of the cytoplasm containing a nucleus with a nuclear pole (NP). Note the lucent area due to the separation of the inner (inm) and outer (onm) nuclear membranes. (c) Low power image of a mid-stage oocyst showing nuclear swelling (N) and increased numbers of lucent cytoplasmic vacuoles (V). (d) Detail part of the cytoplasm showing a swollen nucleus (N), membrane bound lucent vacuoles (V) and mitochondria (Mi) with vesicles embedded in electron dense material. (e) Low power image of a late stage oocyst with abnormal nuclei (N) and the cytoplasm filled with electron lucent vacuoles. (f) Detail from the central region of (e) showing the peripheral location of electron dense chromatin (Ch) typical of apoptotic changes, while the cytoplasm consists of numerous vacuoles (V). (g) Low power image of an early oocyst (end of growth phase) in which the cytoplasm completely fills the oocyst and contains many nuclear profiles (N). (h) Detail of the peripheral cytoplasm limited by the plasmalemma (P) containing mitochondria (M) and nuclei (N). OW – oocyst wall. (i) Mid stage oocyst showing the surface formation of numerous sporozoites (Sp). N – nucleus. (j) Detail showing partially formed sporozoites (Sp) budding from the surface of the cytoplasmic mass. N – nucleus, R – rhoptry. (k) Mature oocysts containing large number of fully form sporozoites (Sp). (l) Detail of cross sections through mature sporozoites (Sp) containing rhoptries (R) and micronemes (MN).

## Discussion

In this study, we examined the location and function of *Plasmodium* DEH using *in silico*, genetic manipulation and cell biological techniques. Lipid metabolism is essential for cellular function, and includes critical pathways for FA synthesis and elongation. DEH is a 3-hydroxyacyl-CoA dehydratase involved in VLCFA synthesis, which interacts with several elongase units, is located at the ER (Beaudoin *et al.*, 2009, Morineau *et al.*, 2016) and has an essential role during development, differentiation, and maintenance of a number of tissue types (Li *et al.*, 2000, Bellec *et al.*, 2002, Pele *et al.*, 2005).

In our previous protein phosphatome study, a putative, catalytically inactive, PTP-like protein with an essential role during sporogony was identified (Guttery *et al.*, 2014), which had been classified as a putative PTPLA by others (Andreeva *et al.*, 2008, Wilkes *et al.*, 2008, Pandey *et al.*, 2014) based on high sequence similarity and e-score values. However, a recent genome-wide functional screen in *P. berghei* showed that PbPTPLA has an essential role in lipid metabolism, specifically during the ELO pathway as a 3-hydroxyacyl dehydratase (DEH) (Stanway *et al.*, 2019). The specific criterion for a PTP-like protein is the presence of a CXXGXXP motif (i.e. the CXXGXXR motif of PTPs, but with the arginine replaced by proline). However, we show here that this motif is not present in either *P. falciparum* or *P. berghei* protein and this, along with its proven function in lipid metabolism (Stanway *et al.*, 2019), suggests that the classification as a phosphatase-like protein is erroneous. Our *in silico* interactome analysis suggests that PfDEH interacts with a number of proteins involved in lipid metabolism, confirming previous functional findings (Stanway *et al.*, 2019) and adding further weight to its annotation as a 3-hydroxyacyl-CoA dehydratase.

Studies in mammalian systems have suggested that the ER-bound DEH catalyzes the third of four reactions in the long-chain FA elongation cycle (Ikeda *et al.*, 2008). Our detailed GFP-based localization analyses showed that the protein is expressed strongly throughout all life-cycle stages analyzed, with protein expression at localized areas in the cytoplasm and as a circular ring-like structure around the nucleus. *In silico* analysis using Predotar and microscopy-based co-localization using ER tracker confirmed the ER location, consistent with a previous study suggesting a role in FAS in *Plasmodium* (Stanway *et al.*, 2019). Phenotypic analysis of DEH function throughout the life cycle confirmed the results of our previous study (Guttery *et al.*, 2014), highlighting that it is essential for oocyst maturation and sporozoite development, but dispensable for asexual blood stage development (Bushell *et al.*, 2017). Time-course analysis at days 7, 14 and 21 after mosquito infection showed that while early-stage Δ*deh* oocysts were comparable in size to WT oocysts, they begin to degenerate at an early stage of development, with a significant decrease in GFP-expressing oocysts even at day 7 post-infection, and as seen previously in other FAE-critical mutant parasites (Stanway *et al.*, 2019). Ultrastructure analysis confirmed that at 14 days post-infection, Δ*deh* oocysts were at an advanced state of degeneration, with no evidence of sporozoite development. Retraction of the oocyst plasmalemma (the parasite plasma membrane) from the oocyst capsule is a crucial first stage in sporozoite development, where sporoblast formation is followed by thousands of sporozoites budding off into the space delineated by the capsule (Aly *et al.*, 2009). A model of this process has been detailed in (Burda *et al.*, 2017). Our study suggests that initiation of mitosis, which results in sporozoite development, does not even commence in Δ*deh* oocysts, since retraction of the plasmalemma and initiation of daughter IMC formation is ablated. The phenotype is similar to that of a cyclin-3 mutant (Roques *et al.*, 2015), with defects leading to abnormalities in membrane formation, vacuolation and subsequent cell death during the later stages of sporogony. However, in contrast to the cyclin-3 mutant, oocyst growth was not affected but sporogony was completely ablated in Δ*deh* parasites, and no transmission was observed in bite-back experiments. This suggests that the parasite cannot scavenge VLCFA from its mosquito host environment, and that DEH (and therefore the ELO pathway) is critical for oocyst mitotic maturation and differentiation. The cells were unable to progress further to form additional lobes and start sporozoite budding in its absence, although the oocyst size was not grossly affected, suggesting that two independent processes drive oocyst formation and sporogony, respectively.

While FASII activity is exclusively in the apicoplast (Shears *et al.*, 2015), our study showed that DEH-GFP is located at the ER, suggesting that it is an active component of the ELO pathway. The genes involved in the ELO pathway include members of the ELO family (1, 2 and 3), of which *ELO2* and *ELO3* are involved in keto- and enoyl-reduction (Kohlwein *et al.*, 2001). In yeast, the dehydratase step is carried out by the DEH-like homologue, Phs1, which has also been characterized as a cell cycle protein with mutants defective in the G2/M phase (Yu *et al.*, 2006). Gene knockout studies for any ELO proteins are few, with a single genome-wide functional analysis showing that the *P. berghei* homologue of PF3D7_0605900 (a putative long chain polyunsaturated fatty acid elongation enzyme) is dispensable during the asexual blood stages (Bushell *et al.*, 2017). In addition, a comprehensive analysis of FAE in *Plasmodium* (Stanway *et al.*, 2019) showed that mutants of a ketoacyl-CoA reductase (KCR) have an identical phenotype to our DEH gene knockout lines, with normal development of ookinetes and oocysts gradually disappearing over the course of development, resulting in the complete ablation of sporogony. In contrast, ELO-A (stage 1 of VLCFA synthesis) mutants were critical for liver stage development. This suggests that reduction of ketoacyl-CoA to hydroxyacyl-CoA and subsequent dehydration of hydroxyacyl-CoA to enoyl-CoA (i.e. stages 2 and 3 of VLCFA synthesis) are the most crucial stages for oocyst maturation and sporogony; whereas the lack of a phenotype during sporogony of the ELO-A deletion may suggest functional redundancy and/or a compensatory mechanism such as an overlapping specificity for condensation of malonyl-CoA by either ELO-B or ELO-C, as suggested in *Trypanosoma brucei* (Lee *et al.*, 2006) and *Toxoplasma gondii* (Ramakrishnan *et al.*, 2012).

In conclusion, our PbDEH analysis using various *in silico, in vitro and in vivo* approaches provides important insights into the crucial role DEH plays during VLCFA synthesis, and how disruption of the gene can affect parasite development in the mosquito. Future studies will elucidate further how lipid metabolism in *Plasmodium* can be explored as a viable target for therapeutic intervention.

## Methods

### Ethics statement

All animal work was performed following ethical approval and was carried out under United Kingdom Home Office Project Licence 40/3344, in accordance with the UK ‘Animals (Scientific Procedures) Act 1986’ and in compliance with ‘European Directive 86/609/EEC’ for the protection of animals used for experimental purposes. Six-to eight-week old female Tuck-Ordinary (TO) (Harlan) or Swiss Webster (Charles River) outbred mice were used for all experiments.

### Identification of conserved domains and evolutionary lineage

The deduced amino acid sequence of PBANKA_134650 (PbPTPLA) now classified as DEH in the manuscript, was retrieved from PlasmoDB (release 27) (Aurrecoechea *et al.*, 2009). Conserved domains in PbPTPLA (DEH) were identified using the Conserved Domain Database (CDD) (Marchler-Bauer *et al.*, 2011), the Simple Modular Architecture Research Tool (SMART) (Schultz *et al.*, 1998) and Protein Family Database (PFAM) (Finn *et al.*, 2008). The deduced amino acid sequence and individual conserved domains were used as BLAST (BLASTP) queries to identify orthologues in PlasmoDB and NCBI protein databases. OrthoMCL database (version 5) was used to identify and retrieve *P. berghei* orthologues (Table S1) (Li *et al.*, 2003). Multiple sequence alignment was performed for the retrieved sequences using ClustalW (Larkin *et al.*, 2007). ClustalW alignment parameters included gap opening penalty (GOP) 10 and gap extension penalty (GOE) 0.1 for the pairwise sequence alignment; GOP 10 and GOE 0.2 was used for multiple sequence alignment. A gap separation distance cut-off of 4 and Gonnet protein weight matrix was used for the alignments. Residue-specific penalty and hydrophobic penalties were used, whereas end gap separation and negative matrix were excluded in the ClustalW alignments. The phylogenetic tree was inferred using the neighbor-joining method, computing the evolutionary distances using the Jones-Taylor-Thornton (JTT) model for amino acid substitution with the Molecular Evolutionary Genetics Analysis software (MEGA 6.0) (Tamura *et al.*, 2013). Gaps and missing data were treated using a partial deletion method with 95% site-coverage cut-off and 1000 bootstrap replicates to generate a phylogenetic tree. iTOL was used to visualize the phylogenetic tree (Letunic *et al.*, 2019). For structure analyses, the secondary structure of the PbDEH was evaluated using PSIPRED (Buchan *et al.*, 2013). I-TASSER, an Iterative threading assembly-based tool, was used to generate 3D structure of PbDEH (Zhang, 2008). The STRING database was used to identify PTPLA interacting proteins (Franceschini *et al.*). For STRING search, we used a cut-off of 0.70 for the parameters of neighborhood, gene fusion, co-occurrence, co-expression, experiments, databases and text mining results. Predotar (Small *et al.*, 2004) was used for inferring Pf and PbDEH subcellular localization.

### Generation of transgenic parasites and genotype analysis

Details of GFP-tagged PTPLA (termed DEH-GFP in this study) and *deh* (PBANKA_1346500) knockout (KO) parasite lines (Δ*deh* in this study) are given in (Guttery *et al.*, 2014). For this study, the KO construct was transfected into the GFPCON wild-type line (Janse *et al.*, 2006), with 3 clones produced by serial dilution.

### Parasite development in the mosquito

*Anopheles stephensi* mosquitoes (3–6 days old) were allowed to feed on anaesthetized mice infected with either wild type or mutant parasites at comparable gametocytemia as assessed by blood smears. Mosquitoes were dissected post-blood meal, on the days indicated. For midgut and salivary gland sporozoites, organs from 10-20 mosquitoes were pooled and homogenized, and released sporozoites were counted using a haemocytometer. For oocyst counts, midguts taken at day 7, 14 and 21 post-infection were harvested, mounted on a slide and oocysts counted using phase or fluorescence microscopy. To quantify sporozoites per oocyst (the ratio of number of sporozoites to number of oocysts), an equal number of mosquitoes from the same cage were used to count the number of oocysts and sporozoites. This number varied among experiments but at least 20 mosquitoes were used for each count. For light microscopy analysis of developing oocysts, at least 20 midguts were dissected from mosquitoes on the indicated days and mounted under Vaseline-rimmed cover slips. ER tracker (ThermoFisher) was used to perform co-localization studies according to manufacturer’s instructions. Images were collected with an AxioCam ICc1 digital camera fitted to a Zeiss AxioImager M2 microscope using a 63x oil immersion objective. Statistical analyses were performed using Graphpad prism software.

### Electron Microscopy

The guts from mosquitoes harvested at 10, 14 and 21 days post-infection were dissected and fixed in 2.5% glutaraldehyde in 0.1 M phosphate buffer and processed for electron microscopy (Guttery *et al.*, 2012). For quantitation, between 10 and 20 oocysts were examined for each group.

### Subcellular fractionation of parasite lysates and detection of DEH

Immunoprecipitation and subcellular fractionation of lysates containing GFP tagged DEH was performed as described previously (Guttery *et al.*, 2014). WT-GFP was used as the control protein in all experiments. In summary, cells from mouse blood infected with the DEH-GFP-expressing parasite were pelleted and then lysed in hypotonic buffer (10 mM Tris-HCl pH 8.4, 5 mM EDTA) containing protease inhibitors (Roche), freeze/thawed twice, incubated for 1 hr at 4°C and then centrifuged at 100,000 g for 30 min. The supernatant was collected as the soluble protein fraction (cytosol). The pellet was resuspended and washed in carbonate solution (0.1M Na_2_CO_3_, pH 11.0) containing protease inhibitors (Roche), and after incubation for 30 min at 4°C the sample was centrifuged again at 100,000 g for 30 min. The supernatant was saved as the peripheral membrane protein fraction (PMF). The residual pellet was solubilized in 4% SDS and 0.5% Triton X-100 in PBS, to form the integral membrane protein fraction (IMF). Samples from these three fractions, containing equal amounts of protein, were then analyzed by western blot using anti-GFP antibody.

## Acknowledgements

We thank Julie Rodgers for maintaining the insectary breeding colony, and Ms. Jessica Stock for assistance in western blot analysis.

## Author contributions

Conceived and designed the experiments: RT, DSG, AAH. Performed the experiments: RT, RJW, RP, DB, DJPF and DSG. Analyzed the data: RT, AAH, RP, RJW, DJPF and DSG. Contributed reagents/materials/analysis tools: DJPF, AAH and RT. Wrote the paper: RP, DSG, RJW, AAH and RT. Performed the functional and GFP tagging experiments: RP, RJW, DB, DSG and RT. Phylogenetics analysis: RP and DSG. RP and DG performed database searches, sequence-based analysis, and other bioinformatics analysis. Electron microscopy experiment: DJPF.

---

**Figure S1: Schematic of the Fatty Acid Synthase II (FASII) and Fatty Acid Elongation (ELO) pathways**

The FASII pathway is localized to the apicoplast (green), whereby ketoacyl-ACP is synthesized by condensation from CoA-activated precursors. The molecule is sequentially reduced, dehydrated and reduced to produce a fatty acyl chain. This cyclical process generates free fatty acids as the final products. The fatty acid elongation (ELO) pathway proceeds on the cytoplasmic face of the endoplasmic reticulum, and catalyzes condensation a malonyl-CoA molecule with a fatty acyl-CoA derived from the FASII pathway. The products of ELO-A, ELO-B and ELO-C are then cyclically reduced, dehydrated and reduced by ketoacyl-CoA reductase (KCR), hydroxyacyl-CoA dehydratase (DEH) and enoyl-CoA reductase (ECR) to finally produce free fatty acids with 2 additional carbons with each cycle.

**Figure S2: Phylogenetic analysis of DEH homologues across different species**

Phylogenetic analysis for PbDEH/PTPLA orthologs was performed using the neighbor joining method with MEGA6 software. Genome wide DEH/PTPLA analysis shows its presence in all organisms studied, including species from all major eukaryotic phyla. The analysis shows the clustering of organisms based on their evolutionary relatedness. Blue – Chordata; Orange – Non-chordata; Purple – Nematoda; Pink – Ameobozoa; Red – Apicomplexa, Euglenozoa and Metamonada; Green – Plants and Algae; Yellow - Fungi and Black - Others.

**Figure S3: ClustalW alignment of *Plasmodium* DEH with human, mouse and other apicomplexan homologues**

ClustalW alignment of *Homo sapiens* (Hs), *Mus musculus* (Ms), *Plasmodium* (Pf, Pb), *Toxoplasma* (Tg) and *Cyclospora* (Cc) homologues of DEH. For apicomplexans, the proteins are currently annotated as PTPLA. Highlighted in red is the CXXGXXP motif that defines PTP-like proteins.

**Figure S4: Predicted 2D and 3D structures of *Plasmodium* DEH**

(A) Secondary structure of *Plasmodium* DEH. (B) Left: Tertiary cartoon view of PbDEH showing the presence of six major hydrophobic membrane-spanning helices followed by coils and an absence of beta sheets, confirming the secondary structure. Right: Tertiary cartoon with surface view of PbDEH.

**Figure S5: Predicted interacting partners of PfDEH**

STRING analysis of PfDEH for potential interacting partners. The table on the right gives the annotation and combined score.

**Figure S6: Representative examples of oocyst degeneration in the mosquito**

Oocysts (63x magnification) in Δ*deh* and WT lines. DIC and GFP images at 7, 10, 14 and 21 dpi. Scale bar = 20 μm.

**Table S1:** Organisms and protein sequences used for phylogenetic analysis.

**Table S2:** Quantitative data related to Figure 3B and 3C.

## References

Aly, A.S.I., et al. (2009). Malaria Parasite Development in the Mosquito and Infection of the Mammalian Host. Annual review of microbiology 63, 195–221.

Andreeva, A.V., et al. (2008). Protozoan protein tyrosine phosphatases. Int J Parasitol 38, 1279–1295.

Aurrecoechea, C., et al. (2009). PlasmoDB: a functional genomic database for malaria parasites. Nucleic acids research 37, D539–543.

Bach, L., et al. (2008). The very-long-chain hydroxy fatty acyl-CoA dehydratase PASTICCINO2 is essential and limiting for plant development. Proc Natl Acad Sci U S A 105, 14727–14731.

Beaudoin, F., et al. (2009). Functional characterization of the Arabidopsis beta-ketoacyl-coenzyme A reductase candidates of the fatty acid elongase. Plant Physiol 150, 1174–1191.

Bellec, Y., et al. (2002). Pasticcino2 is a protein tyrosine phosphatase-like involved in cell proliferation and differentiation in Arabidopsis. Plant J 32, 713–722.

Bisanz, C., et al. (2006). Toxoplasma gondii acyl-lipid metabolism: de novo synthesis from apicoplast-generated fatty acids versus scavenging of host cell precursors. Biochem J 394, 197–205.

Buchan, D.W., et al. (2013). Scalable web services for the PSIPRED Protein Analysis Workbench. Nucleic acids research 41, W349–357.

Burda, P.C., et al. (2017). A Plasmodium plasma membrane reporter reveals membrane dynamics by live-cell microscopy. Sci Rep 7, 9740.

Bushell, E., et al. (2017). Functional Profiling of a Plasmodium Genome Reveals an Abundance of Essential Genes. Cell 170, 260–272 e268.

Costa, G., et al. (2018). Non-competitive resource exploitation within mosquito shapes within-host malaria infectivity and virulence. Nat Commun 9, 3474.

Finn, R.D., et al. (2008). The Pfam protein families database. Nucleic Acids Res 36, D281–288.

Franceschini, A., et al. STRING v9.1: protein-protein interaction networks, with increased coverage and integration. Nucleic Acids Res 41, D808–815.

Guttery, D.S., et al. (2012). A Putative Homologue of CDC20/CDH1 in the Malaria Parasite Is Essential for Male Gamete Development. PLoS Pathog 8, e1002554.

Guttery David S., et al. (2014). Genome-wide Functional Analysis of Plasmodium Protein Phosphatases Reveals Key Regulators of Parasite Development and Differentiation. Cell Host & Microbe 16, 128–140.

Ikeda, M., et al. (2008). Characterization of four mammalian 3-hydroxyacyl-CoA dehydratases involved in very long-chain fatty acid synthesis. FEBS Lett 582, 2435–2440.

Janse, C.J., et al. (2006). High efficiency transfection of Plasmodium berghei facilitates novel selection procedures. Molecular and biochemical parasitology 145, 60–70.

Kihara, A. (2012). Very long-chain fatty acids: elongation, physiology and related disorders. J Biochem 152, 387–395.

Kohlwein, S.D., et al. (2001). Tsc13p is required for fatty acid elongation and localizes to a novel structure at the nuclear-vacuolar interface in Saccharomyces cerevisiae. Mol Cell Biol 21, 109–125.

Larkin, M.A., et al. (2007). Clustal W and Clustal X version 2.0. Bioinformatics 23, 2947–2948.

Lee, S.H., et al. (2006). Fatty acid synthesis by elongases in trypanosomes. Cell 126, 691–699.

Letunic, I., et al. (2019). Interactive Tree Of Life (iTOL) v4: recent updates and new developments. Nucleic acids research 47, W256–W259.

Li, D., et al. (2000). Human protein tyrosine phosphatase-like gene: expression profile, genomic structure, and mutation analysis in families with ARVD. Gene 256, 237–243.

Li, L., et al. (2003). OrthoMCL: identification of ortholog groups for eukaryotic genomes. Genome Res 13, 2178–2189.

Lin, X., et al. (2012). Protein tyrosine phosphatase-like A regulates myoblast proliferation and differentiation through MyoG and the cell cycling signaling pathway. Mol Cell Biol 32, 297–308.

Marchler-Bauer, A., et al. (2011). CDD: a Conserved Domain Database for the functional annotation of proteins. Nucleic Acids Res 39, D225–229.

Mazumdar, J., et al. (2007). Make it or take it: fatty acid metabolism of apicomplexan parasites. Eukaryot Cell 6, 1727–1735.

Morineau, C., et al. (2016). Dual Fatty Acid Elongase Complex Interactions in Arabidopsis. PloS one 11, e0160631.

Pandey, R., et al. (2014). Genome wide in silico analysis of Plasmodium falciparum phosphatome. BMC Genomics 15, 1024.

Pele, M., et al. (2005). SINE exonic insertion in the PTPLA gene leads to multiple splicing defects and segregates with the autosomal recessive centronuclear myopathy in dogs. Hum Mol Genet 14, 1417–1427.

Ramakrishnan, S., et al. (2012). Apicoplast and endoplasmic reticulum cooperate in fatty acid biosynthesis in apicomplexan parasite Toxoplasma gondii. J Biol Chem 287, 4957–4971.

Ramakrishnan, S., et al. (2013). Lipid synthesis in protozoan parasites: a comparison between kinetoplastids and apicomplexans. Prog Lipid Res 52, 488–512.

Roques, M., et al. (2015). Plasmodium P-Type Cyclin CYC3 Modulates Endomitotic Growth during Oocyst Development in Mosquitoes. PLoS Pathog 11, e1005273.

Schultz, J., et al. (1998). SMART, a simple modular architecture research tool: identification of signaling domains. Proc Natl Acad Sci U S A 95, 5857–5864.

Shears, M.J., et al. (2015). Fatty acid metabolism in the Plasmodium apicoplast: Drugs, doubts and knockouts. Molecular and biochemical parasitology 199, 34–50.

Small, I., et al. (2004). Predotar: A tool for rapidly screening proteomes for N-terminal targeting sequences. Proteomics 4, 1581–1590.

Stanway, R.R., et al. (2019). Genome-Scale Identification of Essential Metabolic Processes for Targeting the Plasmodium Liver Stage. Cell 179, 1112–1128 e1126.

Tamura, K., et al. (2013). MEGA6: Molecular Evolutionary Genetics Analysis version 6.0. Mol Biol Evol 30, 2725–2729.

Tarun, A.S., et al. (2009). Redefining the role of de novo fatty acid synthesis in Plasmodium parasites. Trends Parasitol 25, 545–550.

Tehlivets, O., et al. (2007). Fatty acid synthesis and elongation in yeast. Biochim Biophys Acta 1771, 255–270.

van Schaijk, B.C., et al. (2014). Type II fatty acid biosynthesis is essential for Plasmodium falciparum sporozoite development in the midgut of Anopheles mosquitoes. Eukaryot Cell 13, 550–559.

Vaughan, A.M., et al. (2009). Type II fatty acid synthesis is essential only for malaria parasite late liver stage development. Cell Microbiol 11, 506–520.

Wilkes, J.M., et al. (2008). The protein-phosphatome of the human malaria parasite Plasmodium falciparum. BMC Genomics 9, 412.

WHO (2018) World Malaria Report 2018. World Health Organization, pp. 166.

Yu, L., et al. (2006). A survey of essential gene function in the yeast cell division cycle. Mol Biol Cell 17, 4736–4747.

Yu, M., et al. (2008). The fatty acid biosynthesis enzyme FabI plays a key role in the development of liver-stage malarial parasites. Cell Host Microbe 4, 567–578.

Zhang, Y. (2008). I-TASSER server for protein 3D structure prediction. BMC bioinformatics 9, 40.

