## Supplementary figures and images for "*Plasmodium* DEH is ER-localized and crucial for oocyst mitotic division during malaria transmission"

### Figure S1

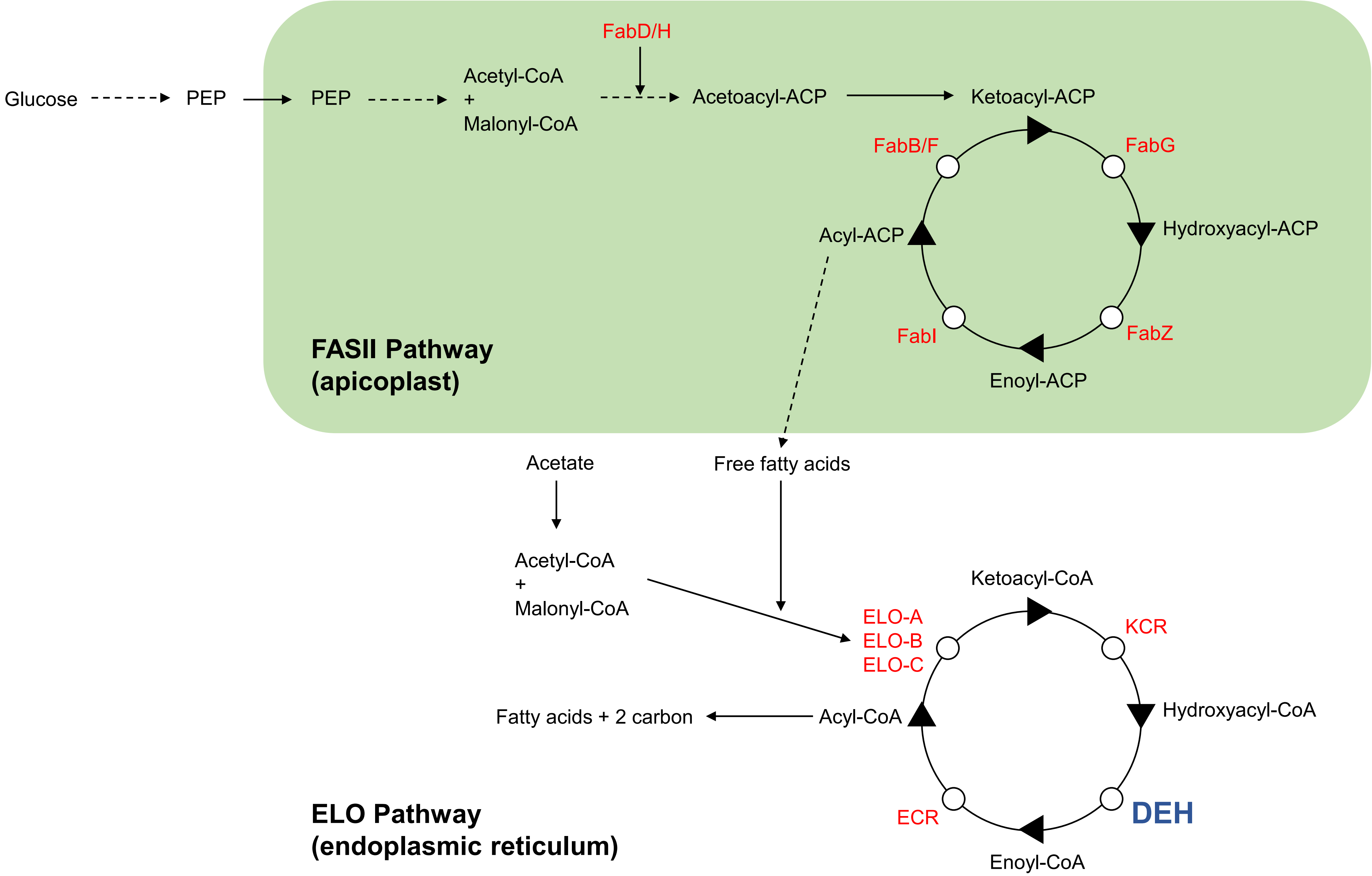

### Figure S2

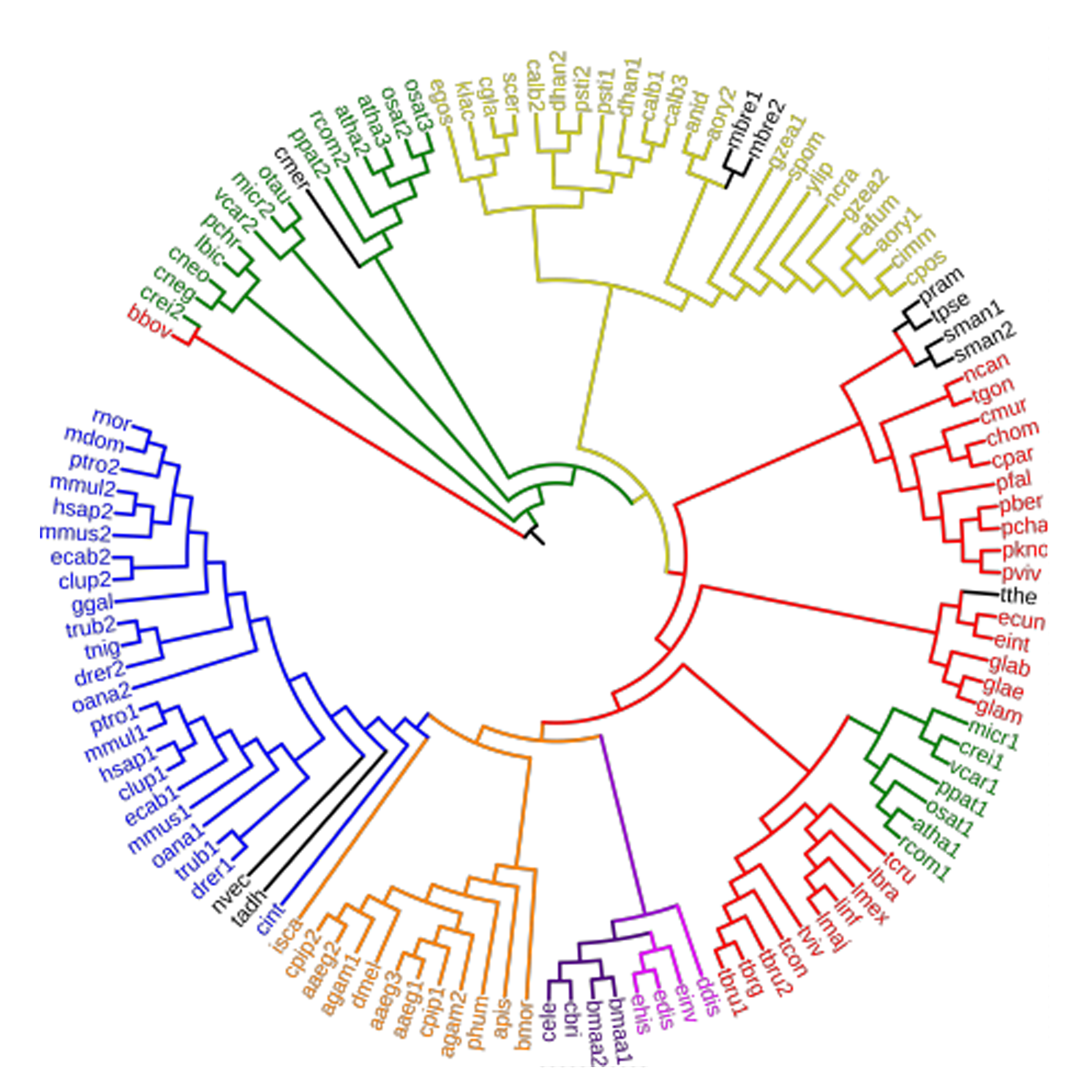

### Figure S4

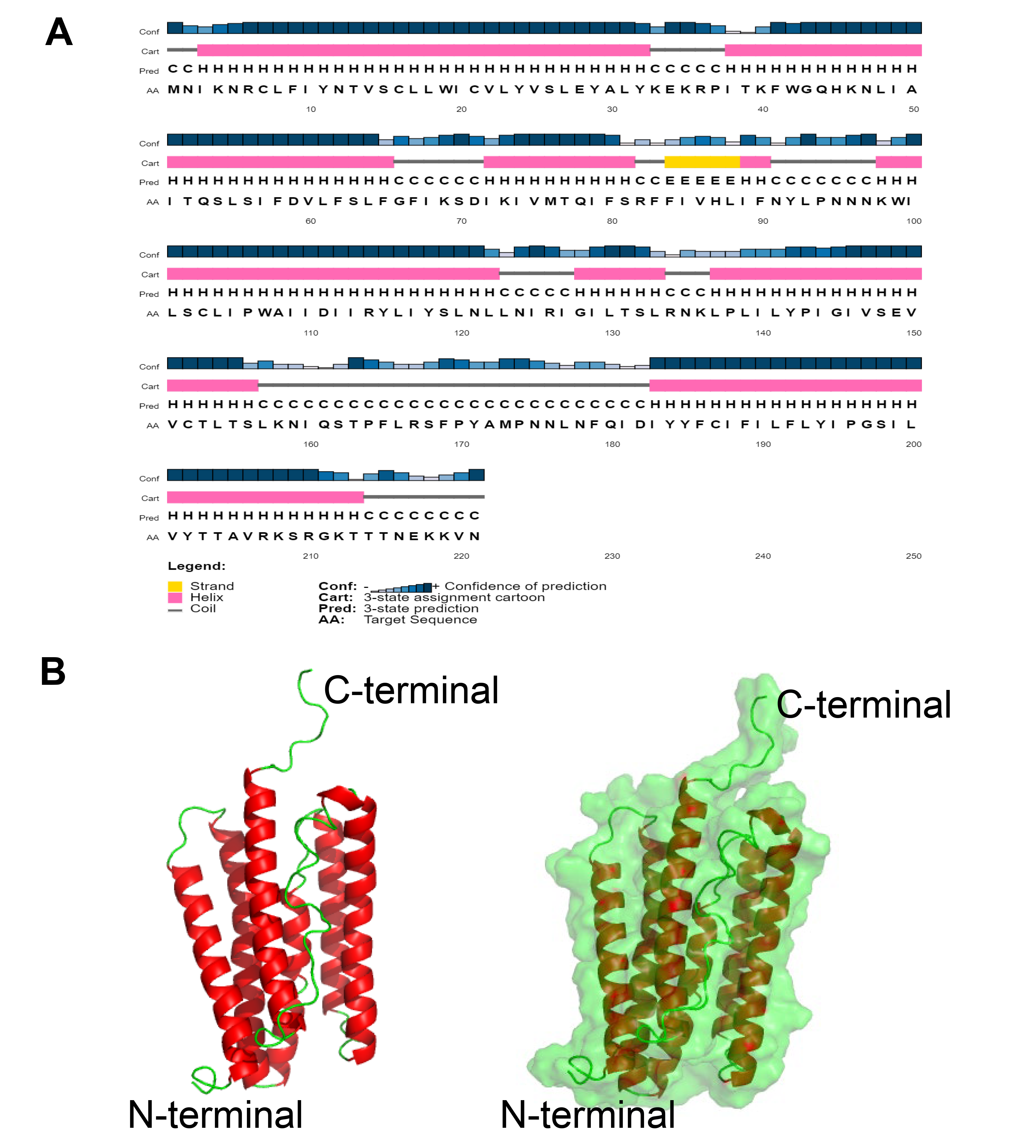

### Figure S5

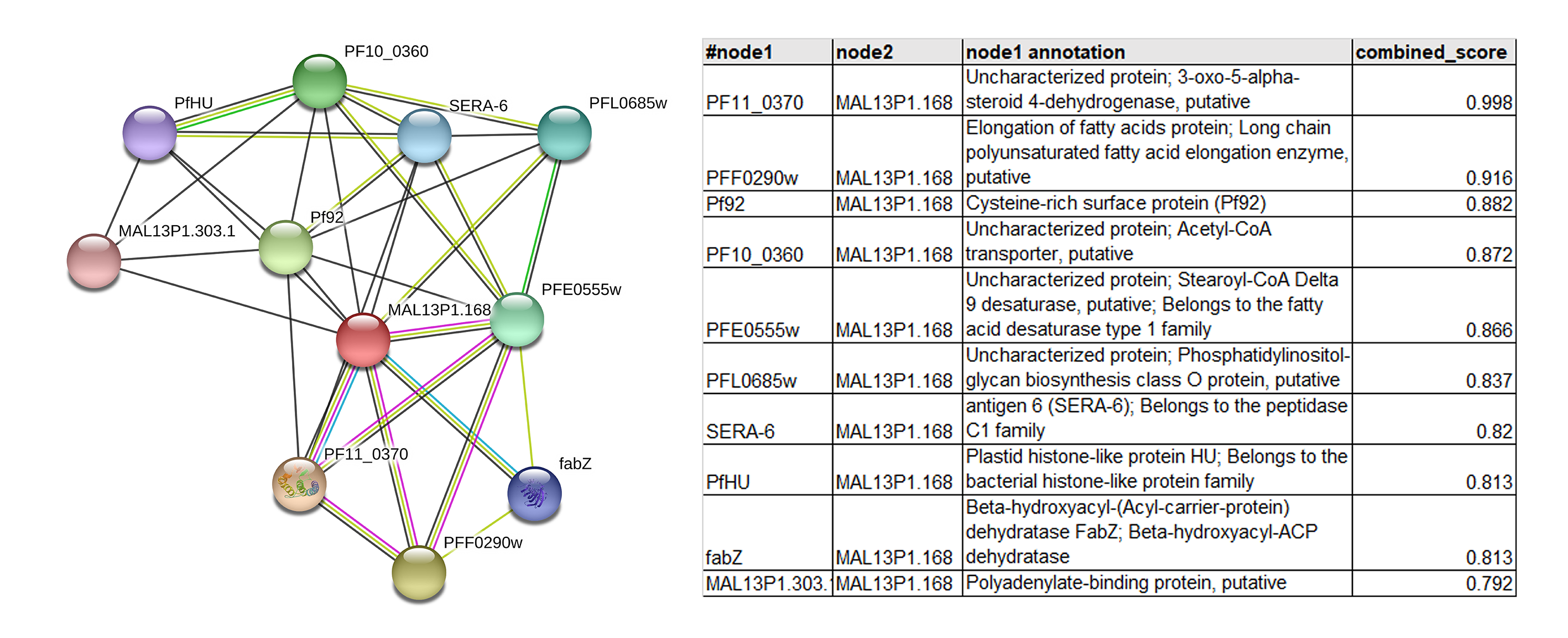

### Figure S6

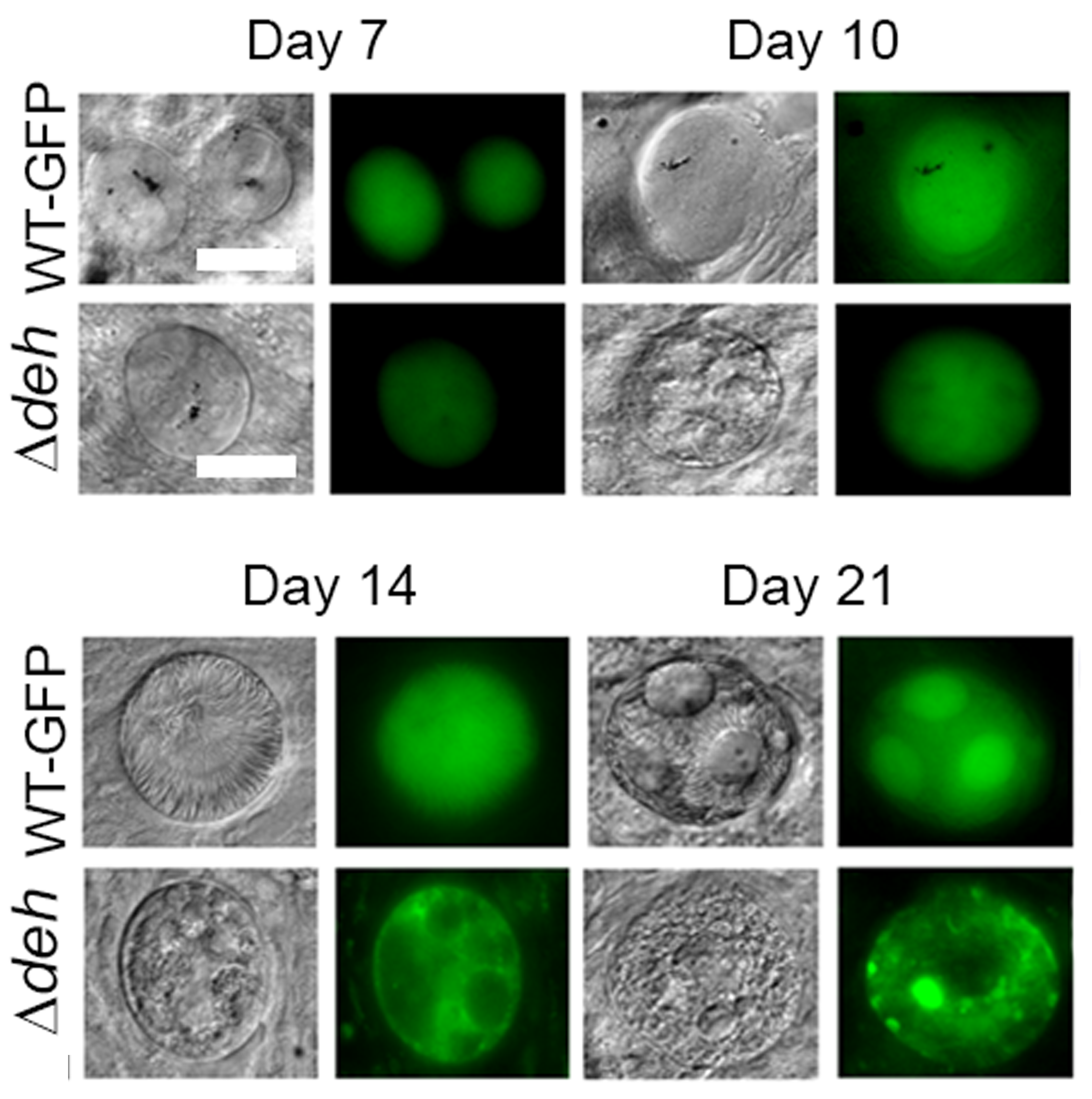
