## Supplementary material for "*Plasmodium* DEH is ER-localized and crucial for oocyst mitotic division during malaria transmission": Figure S3

```
HsHACD2_Q6Y1H2       --------------------------------------MAAVAATAAAKGNGGGGGRAGA 22
MmHACD2_Q9D3B1       --------------------------------------MAAAAATAATKGNGGGSGRVGA 22
HsHACD1_B0YJ81       MGRLTEAAAAGSGSRAAGWAGSPPT---LLPLSPTSPRCAATMASSDEDGTNGGASEA-G 56
MmHACD1_Q9QY80       ---------MGKGDWRQGRVEMPCAHVSRLHKTCVQVRVRVTMASSEEDGTNG-ASEA-S 49
PfPTPLA_PF3D7_133160 ------------------------------------------------------------ 0
PbPTPLA_PBANKA_13465 ------------------------------------------------------------ 0
TgPTPLA_TGARI_311290 ------------------------------------------MAE----AAPARASLAG- 13
CcPTPLA_cyc_02534    ------------------------------------------MAGCASGEMEATSARTG- 17

HsHACD2_Q6Y1H2       GDASGTRKKKG-PGPLATAYLVIYNVVMTAGWLVIAVGLVRAYLAKG-----SYHSLYYS 76
MmHACD2_Q9D3B1       GDSSGARKKKG-PGPVATAYLVIYNVVMTAGWLVIAVGLVRAYLAKG-----SYHSLYYS 76
HsHACD1_B0YJ81       EDREAPGERRR-LGVLATAWLTFYDIAMTAGWLVLAIAMVRFYMEKG-----THRGLYKS 110
MmHACD1_Q9QY80       DEKEAAGKRRR-LGLLATAWLTFYNIAMTAGWLVLAIAMVRFYMEKG-----THRGLYKS 103
PfPTPLA_PF3D7_133160 -------------MSATSKCMLIYNLVCCCLWTVILFTSLQYVFNKE---KYPITTFWSN 44
PbPTPLA_PBANKA_13465 -------------MNIKNRCLFIYNTVSCLLWICVLYVSLEYALYKE---KRPITKFWGQ 44
TgPTPLA_TGARI_311290 ---KVPGPRPP--SLAVHLYLFLYNCVATAAWSVVFFLFAQHVCQRASWTDFAVPALYRS 68
CcPTPLA_cyc_02534    ---ALPGKNSSTRNVLGCLYMLGFNLLCTYAWGVVFLALIRHFVQHSLPGAF-FANAWPE 73

                                        CXXGXXP
HsHACD2_Q6Y1H2       IEKPLKFFQTGALLEILHCAIGIVPSSVVLTSFQVMSRVFLIWAVTHSVKEVQSEDSVLL 136
MmHACD2_Q9D3B1       IERPLKFFQTGALLEILHCAIGIVPSSVVLTSFQVMSRVFLIWAVTHSVKEVQSEDSVLL 136
HsHACD1_B0YJ81       IQKTLKFFQTFALLEIVHCLIGIVPTSVIVTGVQVSSRIFMVWLITHSIKPIQNEESVVL 170
MmHACD1_Q9QY80       IQKTLKFFQTFALLEVVHCLIGIVPTSVLVTGVQVSSRIFMVWLITHSIKPIQNEESVVL 163
PfPTPLA_PF3D7_133160 YKNIITITQSLAIFEIFFTIIGIINSVVSIVTIQVFSRLFVVYLIFNFLPNTNKWI--LS 102
PbPTPLA_PBANKA_13465 HKNLIAITQSLSIFDVLFSLFGFIKSDIKIVMTQIFSRFFIVHLIFNYLPNNNKWI--LS 102
TgPTPLA_TGARI_311290 LEFPLLFAQSMQVMEVLHAAAGIVRSGVMTTLTQVFSRLQLVLFLFRVVPVTHENAAFCS 128
CcPTPLA_cyc_02534    LEHPLFFAQSLAILEIGHSL---------------------------------------- 93

HsHACD2_Q6Y1H2       FVIAWTITEIIRYSFYTFSLLNH-------------------------LPYLIKWARYTL 171
MmHACD2_Q9D3B1       FVIAWTITEIIRYSFYTFSLLNH-------------------------LPYIIKWARYTL 171
HsHACD1_B0YJ81       FLVAWTVTEITRYSFYTFSLLDH-------------------------LPYFIKWARYNF 205
MmHACD1_Q9QY80       FLVSWTVTEITRYSFYTFSLLDH-------------------------LPHFIKWARYNL 198
PfPTPLA_PF3D7_133160 CLIAWAIIDIIRYLFYSLNILNLRFN-------------------------ILASLRKKL 137
PbPTPLA_PBANKA_13465 CLIPWAIIDIIRYLIYSLNLLNIRIG-------------------------ILTSLRNKL 137
TgPTPLA_TGARI_311290 LIAAWCLAELLRYPFFCAQELLLCIHHKEAKKAFGDDAATAIVKSKTEAPMILRWLRYSG 188
CcPTPLA_cyc_02534    ---------------------------------LG------SEASPREVPFLLKWLRYSG 114

HsHACD2_Q6Y1H2       FIVLYPMGVSGELLTIYAALPFVRQA---GLYSISLPNKYNFSFDYYAFLILIMISYIPI 228
MmHACD2_Q9D3B1       FIVLYPMGVTGELLTIYAALPFVRQA---GLYSISLPNKYNFSFDYHAFLILIMISYIPL 228
HsHACD1_B0YJ81       FIILYPVGVAGELLTIYAALPHVKKT---GMFSIRLPNKYNVSFDYYYFLLITMASYIPL 262
MmHACD1_Q9QY80       FIILYPVGVAGELLTIYAALPYVKKS---GMFSVRLPNKYNVSFDYYYFLLITMASYIPL 255
PfPTPLA_PF3D7_133160 PLILYPIGITSEIVCTLASLNNIYATPFLRTYPYSMPNNINFQIDIYYFCIVVLILYIPG 197
PbPTPLA_PBANKA_13465 PLILYPIGIVSEVVCTLTSLKNIQSTPFLRSFPYAMPNNLNFQIDIYYFCIFILFLYIPG 197
TgPTPLA_TGARI_311290 FTFLYPVGITSEVVCMLSGLSTLQ-LPSFTHFPAPMPNALNFEVNLHGLYVLLLLTYIPG 247
CcPTPLA_cyc_02534    FTVLYPIGIASEVVCMCSSLSVLRSSPAFAQFPTPMPNKLNFQLSLYWAYILLCLYVPG 174

HsHACD2_Q6Y1H2       FPQLYFHMIHQRRKILSHTEEHKKFE----- 254
MmHACD2_Q9D3B1       FPQLYFHMIHQRRKVLSHTEEHKKFE----- 254
HsHACD1_B0YJ81       FPQLYFHMLRQRRKVLHGEVIVEKDD----- 288
MmHACD1_Q9QY80       FPQLYFHMLRQRRKVLHGEVIAEKDD----- 281
PfPTPLA_PF3D7_133160 SILLYATAVRKSKQKIPIPEKKSDGANKKKI 228
PbPTPLA_PBANKA_13465 SILVYTTAVRKSRGKTTTNEKKVN------- 221
TgPTPLA_TGARI_311290 SFLLYSHMLRQRKKHLYGAGSEEKKTQ---- 274
CcPTPLA_cyc_02534    SVQLYNHMLKQRRKYLYDAGQDTAAKKDK-- 203
```
